# GTShark: Genotype compression in large project

**DOI:** 10.1101/494104

**Authors:** Sebastian Deorowicz, Agnieszka Danek

## Abstract

**Summary:** Nowadays large sequencing projects handle tens of thousands of individuals. The huge files summarizing the findings definitely require compression. We propose a tool able to compress large collections of genotypes as well as single samples in such projects to sizes not achievable to date.

**Availability and Implementation:** https://github.com/refresh-bio/GTShark

**Supplementary information:** Supplementary data are available at publisher’s Web site.

## 1 Introduction

Rapid decrease of genome sequencing costs allows many sequencing projects to grow to impressive sizes. The numbers of individuals covered by the Haplotype Resource Consortium (McCarthy *et al.*, 2016) or Exome Aggregate Consortium (Layer *et al.*, 2016) projects are counted in tens of thousands and even lager projects are on the go.

The aggregate results of such projects are usually stored in the Variant Call Format (VCF) files (Danecek *et al.*, 2011), in which the genome variations occupies successive rows. Each tab-separated row contains nine mandatory fields describing the variant and some (possibly large) number of optional values representing genotypes. The sizes of VCF files are huge, so gzip is a common solution partially resolving the storage and transfer problems. Nevertheless, a lot of effort was made to provide even better compression and sometimes speeding up the accession to the data. The most remarkable attempts were TGC (Deorowicz *et al.*, 2013), PBWT (Durbin, 2014), BGT (Li, 2015), GTC (Danek and Deorowicz, 2018). The first of them focused just on the compression ratio, while the remaining aimed at the rapid queries support with good compression ratios.

In this article, our goal is to provide the best compression ratio for a collection of genotypes. Moreover, the compressed database can serve as a knowledge base, which allows to astonishingly reduce sizes of files of newly sequenced individuals.

Our tool, GTShark, is based on the Positional Burrows–Wheeler Transform (PBWT) introduced in (Durbin, 2014). The key idea of PBWT is to permute a vector of genotypes for each single variant to order the samples according to the genotypes of previous variants. Due to the linkage disequilibrium property the neighbor genotypes (after the permutation) are likely to be the same. Thus, the permuted vector of genotypes is usually composed of a few runs of 1s (presence of alternate value) and 0s (presence of the referential value). PBWT and BGT (Li, 2015) store the run lengths using a simple run encoding scheme. BGT preserves also a permutation of samples each 8192nd variant to allow fast random access queries.

## 2 Methods

In GTShark, we essentially follow the same way. Nevertheless, there are several differences. First, we use a generalized PBWT (designed for non-binary alphabets in (Deorowicz *et al.*, 2018)) to directly support multi-alleles and unknown genotypes. Second, since we aim at the compression ratio, we do not store the intermediate permutations. Third, we employ an entropy coder (namely range coder) with special contextual modeling, which improves the compression ratio more than 2-fold. Fourth, we slightly modify the generalized PBWT, i.e., we resign from permuting vectors of samples for extremely rare (or frequent) variants. The experiments show that this improves the compression ratio by up to 10 percent.

A unique feature of GTShark is the ability to compress external, single-sample, VCF files with a use of the compressed collection as a reference. Such a scenario can appear, for example, in large sequencing projects, like the UK Biobank (Bycroft *et al.*, 2018), in which the genotypes are determined using microarrays, when a set of variants is fixed. To compress such sample we traverse the compressed collection and for each genotype we determine the position in the permuted vector in which this genotype would be placed if the sample were a part of the collection. We make use of the found neighborhood to predict and store the genotype compactly. More details of the algorithms can be found in the Supplementary Material.

The compressor was implemented in the C++14 language and is distributed under GNU GPL 3 licence. It is slightly parallelized: the PBWT making and the remaining parts are executed in separate threads. Thus, the maximal parallelization gain is 2-fold but in practice it is much smaller.

## 3 Results

For evaluation we used two *H.sapiens* datasets: the 1000 Genome Project Phase 3 (1000GP3: 2504 samples, 84.80M variants) and the Haplotype Reference Consortium (HRC: 27,165 samples, 40.40M variants).

The comparison of compression ratios and times of the state-of-the-art methods, i.e., TGC, PBWT, BGT, GTC, and the proposed GTShark is shown in Table 1 and Supplementary Material. GTShark is the clear winner in compression ratio. The running times of PBWT, BGT, GTC, GTShark algorithms are similar as they are dominated by processing of VCF files. GTShark is the fastest in compression due to slight parallelization of the code and some small technical improvements in parsing the VCF files. GTShark allows also to extract a single sample from a compressed collection in an average time about 11.5 minutes.

**Table 1.**
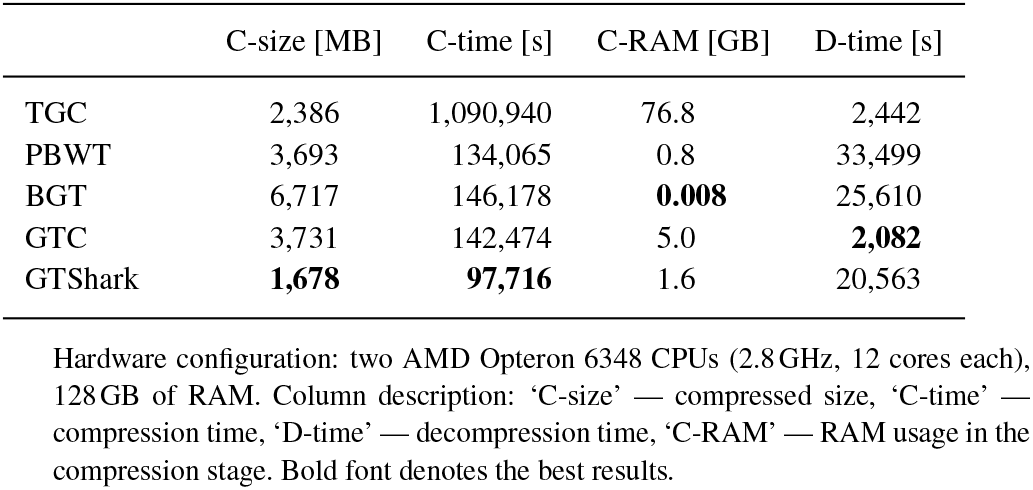
Comparison of compressors of VCF files for HRC collection (40.40 M variants, 27,165 samples, 4310 GB of VCF, 69.7 GB of gzipped VCF).

|  | C-size [MB] | C-time [s] | C-RAM [GB] | D-time [s] |
| --- | --- | --- | --- | --- |
| TGC | 2,386 | 1,090,940 | 76.8 | 2,442 |
| PBWT | 3,693 | 134,065 | 0.8 | 33,499 |
| BGT | 6,717 | 146,178 | **0.008** | 25,610 |
| GTC | 3,731 | 142,474 | 5.0 | **2,082** |
| GTShark | **1,678** | **97,716** | 1.6 | 20,563 |

Hardware configuration: two AMD Opteron 6348 CPUs (2.8GHz, 12 cores each), 128GB of RAM. Column description: 'C-size' — compressed size, 'C-time' — compression time, 'D-time' — decompression time, 'C-RAM' — RAM usage in the compression stage. Bold font denotes the best results.

In the next experiment, we measured the compression of external samples. We used the HRC dataset here and processed as follows. We excluded 100 randomly chosen samples to obtain the collection of 27,065 samples, for which we built the compressed database of total size 1674MB. Then we compressed the excluded samples one by one taking the compressed collection as a reference. On average GTShark processed a single sample in about 12 minutes and compacted it to 65.5 KB. To the best of our knowledge, the most recent experiment of this type was described in (Pavlichin *et al.*, 2013). The authors used a reference human genome and the dbSNP database as a knowledge base. They were able to compress single individual genotypes from the 1000GP Phase 1 (39.7M variants) to about 2.5 MB. The results should not be, however, compared directly as in the Pavlichin et al. experiment the compressed samples contained variants absent from the reference database, and in our experiment we assumed that sets of variants in the sample and reference data are the same. Nevertheless, the comparison shows what is possible in such restricted scenario.

In the final experiment, we investigated the impact of the selected knowledge base. We used Chromosome 11 data from the 1000GP3 containing 2504 samples from 26 populations. Initially, we divided the VCF file into 26 files using the population criteria. The ASW and MXL populations were significantly smaller than the rest and we excluded them from further studies. The remaining populations have cardinalities at least 85 and we randomly subsampled the larger ones to obtain 24 VCF files each containing exactly 85 samples. Then we used GTShark to compress each VCF file obtaining 24 compressed collections serving as references in the rest of the experiment. We also constructed 24 larger referential collections. Each of them, composed of 23 population VCF files and containing 23 × 85 = 1955 samples, was also compressed using GTShark. Then we compressed each single-sample VCF file (i.e., 24 × 85 = 2040 samples in total) using each of the smaller (85-sample) collections as references. The results were averaged over all 85 samples from each population. For easier interpretability of Fig. 1 we grouped the populations in 5 superpopulations (African, American, East Asian, European, and South Asian). The columns are described by the referential population. As one could expect the compression is the best when a dataset from the same superpopulation is used as a reference. The last column (beyond the matrix) shows the results for larger collections (containing all populations except for the one that is compressed).

**Fig. 1.**
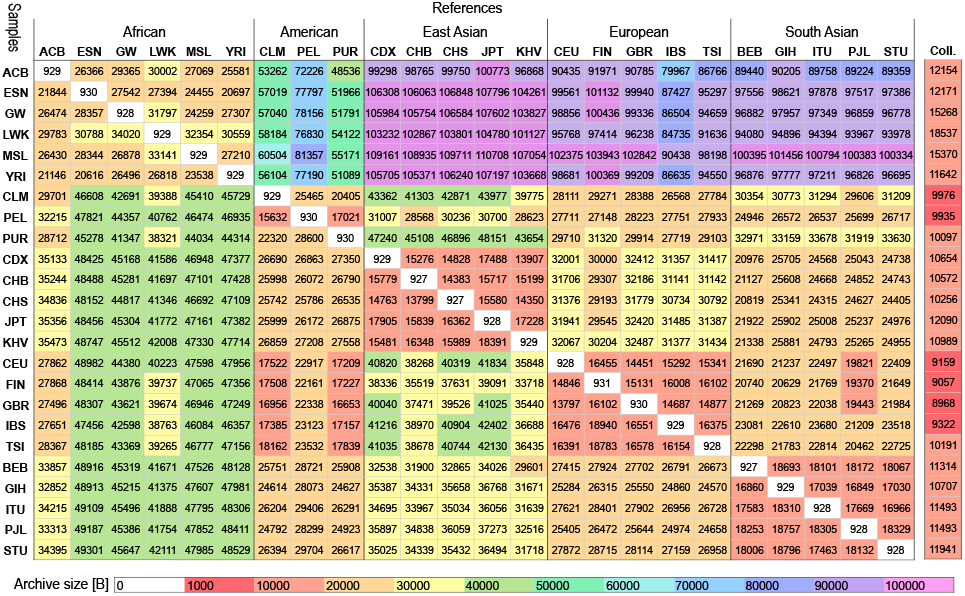
Comparison of single-sample GTShark compression for 24 populations (data: 1000GP3, Chromosome 11), sizes of sample archives. The references with population codes are made up of 85 samples from the population. The “Coll.” reference is a collection of 1955 samples from 23 population (without population matching compressed sample).

## 4 Conclusions

The proposed algorithm compressed large collections of genotype data significantly better than the existing methods and was the fastest. Its unique feature is a mode in which the compressed collection of genotypes serves as a reference for external single-sample data. In this scenario we were able to shrink down the human genome to about 65 KB.

## Supporting information

