## Supplementary material for "GTShark: Genotype compression in large project"

### Supplementary material for article: GTShark: Genotype compression in large project

Sebastian Deorowicz Agnieszka Danek

December 11, 2018

**2 Examined programs 6**

**3 Datasets 7**

**4 Evaluation 8**
4.1 Compression and decompression of a full dataset—compressors comparison . . . 8
4.2 Extraction of a single sample . . . . 9
4.3 Compression and decompression of a single sample using previously compressed dataset as a reference . . . . 9
4.4 Using population as a knowledge base . . . . 10

**5 Environment 11**

**6 Additional results 12**
6.1 Compression and decompression of a full dataset—compressors comparison . . . 12
6.2 Extraction of a single sample . . . . 13
6.3 Compression and decompression of a single sample using previously compressed dataset as a reference . . . . 13
6.4 Using population as a knowledge base . . . . 13

### 1 Algorithmic details

Below we give more details of the algorithms used in GTShark. We focus on the compression stages as the decompression is almost symmetric.

#### 1.1 Input VCF files

Each input VCF file must have the homogeneous ploidy (1 or 2). Therefore, files not satisfying this feature must be split before compression to a few files by external tools.

#### 1.2 Preprocessing of VCF files

The input VCF files can have multiallele variants, i.e., the variants for which more than one alternate value is possible. Such files are handled by our compressor. When the files are loaded the multiallele variants are split into as many variants (for the same position in a genome) as the number of alternate alleles. Therefore, the valid genotypes for each allele are: 0 (the referential value), 1 (the alternate value), 2 (some other alternate value), and 3 (the genotype is unknown). Such approach significantly simplifies handling multiallele variants and was used previously also in [4, 1].

For diploid genomes we internally split the genotypes into two haploid genomes so in the rest of the algorithm we can work in exactly the same way independently on the initial ploidy.

#### 1.3 Positional Burrows–Wheeler Transform for non-binary alphabets

The original PBWT algorithm proposed in [3] was designed for binary alphabets. Recently, it was straightforwardly generalized to non-binary alphabets [2]. In GTShark we use this recent variant (gPBWT) as it directly supports the 4-value alphabet we are working on.

#### 1.4 Special treatment of extremely rare or extremely frequent genotypes

We start processing of each variant from determination of a histogram of values, a necessary step in the gPBWT algorithm. Then, we determine the dominant (the most frequent) value. If the total number of non-dominant values are very small (we use the threshold 10 for the 1000GP3 data and 20 for the HRC data) we do not permute.

#### 1.5 Contextual modelling in the collection compression

After gPBWT the genotype values usually form a sequence of long runs. Formally, the variant genotypes can be represented as a sequence of tuples $\langle v, \ell \rangle$, where $v$ is a genotype value (from 0 to 3) and $\ell$ is the run length. The complete sequence is:

$$\langle v_1, \ell_1 \rangle, \langle v_2, \ell_2 \rangle, \ldots, \langle v_k, \ell_k \rangle.$$

We store this sequence with a use of an entropy coder. In the present implementation a range coder is employed here.

For tuple $\langle v_i, \ell_i \rangle$ we firstly encode the value $v_i$ taking as a context 4 recently processed values, i.e., $(v_{i-1}, v_{i-2}, v_{i-3}, v_{i-4})$ (for completeness we define $v_j = 4$ for $j \leq 0$).

The encoding of the run length $\ell_i$ is as follows. Initially we determine the range $r_i$ of length to which $\ell_i$ belongs. The possible ranges are:

- 0 for $\ell_i = 0$ (it will be explained below when the run length can be 0),
- 1 for $\ell_i = 1$,
- 2 for $2 \leq \ell_i < 4$,

- 3 for $4 \le \ell_i < 8$,
- 4 for $8 \le \ell_i < 16$,
- 5 for $16 \le \ell_i < 32$,
- 6 for $32 \le \ell_i < 64$,
- 7 for $64 \le \ell_i < 128$,
- 8 for $128 \le \ell_i < 256$,
- 9 for $256 \le \ell_i < 512$,
- 10 for $512 \le \ell_i$.

The range $r_i$ is encoded using $(v_i, v_{i-1}, v_{i-2}, v_{i-3})$ as a context.

Ranges 0 or 1 uniquely specify the current run length. For range values from 2 to 9 we encode the value $\ell_i - 2^{r_i - 1}$. The context here is $(v_i, r_i)$. For range 10 we treat the length $\ell_i$ as 24-bit integer and split it into bytes (from the most significant one): $b_2, b_1, b_0$. Then, we encode: $b_2$ with context $(r_i)$; $b_1$ with context $(r_i, b_2)$; $b_0$ with context $(r_i, b_2, b_1)$.

The exceptional run length 0 is used instead of $\ell_k$, i.e., the last tuple in a sequence. We can use such a trick as we know the number of samples in the collection and in decompression we can directly calculate $\ell_k$. Moreover length 0 serves as a flag denoting the end of the variant.

The way the lengths are encoded limits (in the current implementation) the number of samples in an input VCF file to $2^{24} \approx 16.7 \times 10^6$, which seems to be a sufficiently large value. Nevertheless, the support for larger collections is straightforward as it suffice to treat $\ell_i$ as, e.g., 32-bit integer and store 4 bytes of length.

#### 1.6 Compression of external single-sample VCF files

The compression of external single-sample VCF files requires that the sets of variants in both the collection and the external sample are exactly the same. Below, we describe how the genotypes of the successive variants in the external sample are predicted and how these predictions are used in the encoding.

Similarly as for collection of samples, the ploidy in the external sample must be homogeneous (and the same as for the collection). If the sample is diploid it is (internally) split into two haploid ones (but this is not visible to the user).

##### 1.6.1 Virtual placement of external sample in a collection

Let the collection contain $n$ samples. The initial permutation of them is $(0, 1, \ldots, n-1)$. We assume that at the beginning our external sample is at the position $n$. Then we proceed as follows.

For each variant we decode the series of runs:

$$\langle v_1, \ell_1 \rangle, \langle v_2, \ell_2 \rangle, \ldots, \langle v_k, \ell_k \rangle$$

and calculate histogram $H$ of occurrences of values. Then we calculate cumulative histogram $H^*$ defined as

$$H^*[i] = \sum_{j=0}^{i-1} H[j].$$

We also have a virtual position $p$ of the external sample in the permutation for the previous variant, which means that the external sample should be located between samples $p-1$ and $p$. Then, we proceed as shown in Figure 1.

**procedure** EXT-SAMPLE-VARIANT-PLACEMENT($v$, $R$, $H^*$, $p$)

Input: $v$ — *current genotype value*
$R$ — *sequence of runs* $\langle v_1, \ell_1\rangle, \langle v_2, \ell_2\rangle, \ldots, \langle v_k, \ell_k\rangle$
$H^*$ — *cumulative histogram of symbols according to* $R$
$p$ — *position of the external sample after processing previous variant*

Output: $N_{\rm p}$ — *neighbour (predecessor) run*
$N_{\rm s}$ — *neighbour (successor) run*
$p^*$ — *position of the external sample after processing the variant*

```
N_p ← ⟨0,0⟩;    N_s ← ⟨0,0⟩
j ← 0      {current position in vector}
c ← 0      {counter for current genotype value}
if p = 0 then
    N_s ← ⟨v_1, ℓ_1⟩
else
    for i ← 1 to k do
        if p = j + ℓ_i then
            N_p ← ⟨v_i, ℓ_i⟩
            if i + 1 ≤ k then N_s ← ⟨v_{i+1}, ℓ_{i+1}⟩
            if v = v_i then c ← c + ℓ_i
            break
        else if p < j + ℓ_i then
            N_p ← ⟨v_i, p − j⟩
            N_s ← ⟨v_i, ℓ_i − (p − j)⟩
            if v = v_i then c ← c + p − j
            break
        j ← j + ℓ_i
        if v = v_i then c ← c + ℓ_i
if permuting-variant then p* ← H*[v] + c
else p* ← p      {do not permute for extremely rare of frequent variants}
return N_p, N_s, p*
```

Figure 1: Pseudocode of the algorithm finding the location of external sample in a compressed collection of genotypes

Roughly speaking we make gPBWT for the variant and the external-sample genotype but as we do not need the complete permutation we just look for the position of the external-sample genotype. We also return the runs before ($N_{\rm p}$) and after ($N_{\rm s}$).

##### 1.6.2 Context modelling for external-sample genotypes

For encoding of the external-sample genotypes we make use of the runs before and after the virtual position of the genotypes. For example, if the external genotype is located between two runs of 0s it is likely that the external genotype is also 0 and taking such predictions should help in compact representation of the actual genotype value.

More formally, let us denote $N_{\rm p} = \langle v_{\rm p}, \ell_{\rm p}\rangle$ and $N_{\rm s} = \langle v_{\rm s}, \ell_{\rm s}\rangle$. We encode the value of external genotype in a context composed of 6 values. The first two are $v_{\rm p}$ and $v_{\rm s}$. The next two are calculated from run lengths: $\left\lfloor \frac{\lfloor \log \ell_{\rm p} \rfloor + 1}{4} \right\rfloor$ and $\left\lfloor \frac{\lfloor \log \ell_{\rm s} \rfloor + 1}{4} \right\rfloor$. The last two are: $\left\lfloor \frac{\lfloor \log lcs_{\rm p} \rfloor + 3}{4} \right\rfloor$ and $\left\lfloor \frac{\lfloor \log lcs_{\rm s} \rfloor + 3}{4} \right\rfloor$, where $lcs_{\rm p}$ and $lcs_{\rm s}$ are the lengths of the common suffixes between the external sample and neighbor samples from the collection.GTShark

The calculation of the common suffixes between the external sample and the neighbor from the database requires explanation. Let us assume that before processing of the current variant the length of the common suffix between the external sample and neighbor sample at position

$p$ is $x$. If $v = v_s$ then after processing the current variant $x$ will change to $x + 1$. (For technical reasons, in the implementation, this value is upper bounded by 2048.) In the opposite case, we have to calculate $lcs_s$ without decompressing large part of the database. Fortunately, this can be achieved in a similar way as extracting single sample from the compressed collection, which is also implemented in GTShark in the similar manner like in PBWT [3], BGT [4].

#### 1.7 Compression of genotype descriptions

GTShark allows also to compress the variant descriptions, i.e., the nine mandatory fields. To this end we gather the field values (from the complete database) separately into 9 sequences. The position values are delta-encoded. Then each sequence is compressed by the LZMA algorithm.

#### 2 Examined programs

The following programs were used in the experimental part.

- BGT v. 1.0-r283-dirty, `https://github.com/lh3/bgt`
- GTC v. 1.1, `https://github.com/refresh-bio/GTC`
- PBWT, `https://github.com/richarddurbin/pbwt`
- TGC, `https://github.com/refresh-bio/TGC`
- GTShark (the proposed algorithm), `https://github.com/refresh-bio/GTShark`

### 3 Datasets

This section presents datasets used in the experimental part.

Each sample in the VCF files from these datasets is described only by a genotype data, encoded as allele values (`GT` field in a VCF file).

##### The 1000 Genomes Project — Phase 3 (1000GP3)

The 1000 Genome Project data sets were downloaded from:
`ftp://ftp.1000genomes.ebi.ac.uk/vol1/ftp/release/20130502/`

It contains 2,504 samples.

The VCF files for all autosomes and chromosome X contain diploid genotype data for all 2,504 samples, while the VCF file for chromosome Y contains haploid genotype data for 1,233 male samples.

##### The Haplotype Resource Consortium (HRC)

The HRC data sets were downloaded from the European Genom-phenom Archive (Study EGAS00001001710, Dataset EGAD00001002729).

It contains 27,165 samples.

The VCF files for all autosomes and two pseudo-autosomal regions of chromosome X/Y (called X_PAR1 and X_PAR2) contain diploid genotype data for: a) all 27,165 samples in case of chromosomes 2–22, b) 22,691 samples in case of chromosome 1, c) 26,199 samples in case of chromosome X/Y.

The VCF file for non pseudo-autosomal region of chromosome X/Y contain mixed (haploid and diploid) genotype data for 22,691 samples. We split this VCF file into two separate VCF files: X_nonPAR_FEMALE containing diploid genotype data for 14,460 female and X_nonPAR_MALE containing haploid genotype data for 11,739 males.

### 4 Evaluation

This section provides details of the experiments performed.

#### 4.1 Compression and decompression of a full dataset—compressors comparison

In this experiment both datasets, 1000GP3 and HRC, were compressed and decompressed using TGC, BGT, GTC, PBWT and proposed GTShark. Each VCF file was processed separately. The output of decompression was, when possible, redirected to null device.

The exact command line parameters of all programs are given below (input VCF is `input.vcf` and archive name is `archive`).

##### TGC

- Compression:

```
./VCF2VDBV  input.vcf
ls -l *.bv > bv_files_list
./tgc c archive  bv_files_list
```

- Decompression:

```
./tgc d archive
```

Note: The VCF2VDBV tool (part of TGC) was modified in order to parse all VCF variants and their description (original tool copes only with variants from phase 1 of 1000GP). It does produce correct Bit Vectors (BV) for all genotypes (which can be compressed with tgc), but the Variant Database it produces is limited and does not allow to recreate the initial VCF. Hence, the decompression here was only to BV (stored on disk).

##### BGT

- Compression:

```
./bgt import -S -o archive input.vcf.gz
```

- Decompression:

```
./bgt view -b -u archive > /dev/null
```

##### GTC

- Compression:

```
./gtc compress -t 1 -o archive input.vcf.gz
```

- Decompression:

```
./gtc view -b -c 0 archive > /dev/null
```

Note: parameter "`-p 1`" was added for compression of VCF files containing haploid genotype data.

##### PBWT

- Compression:

```
./pbwt  -readVcfGT input.vcf.gz -writeAll archive
```

- Decompression:

```
./pbwt -readAll archive -writeBcf - > /dev/null
```

##### GTShark

- Compression:

```
./gtshark compress-db -nl 10  input.vcf.gz archive
```

- Decompression:

```
./gtshark decompress-db  -b -c 0 archive -  > /dev/null
```

Note: Parameter `nl` was set to 10 for the 1000GP3 dataset and to 20 for the HRC dataset.

#### 4.2 Extraction of a single sample

In this experiment GTShark was used to extract 100 random single samples from the previously compressed HRC and 1000GP3 datasets (archives were created in the first experiment).

The exact command line parameters are given below (compressed dataset is `archive`, sample id is `ID`). The output was redirected to null device.

- Sample extraction:

```
./gtshark extract-sample -b -c 0 archive ID - > /dev/null
```

#### 4.3 Compression and decompression of a single sample using previously compressed dataset as a reference

In this experiment, GTShark was used to compress VCF files describing single samples in reference to a previously compressed dataset.

The reference dataset was made of HRC dataset with 100 random samples (50 males and 50 females) excluded from it (all of these 100 samples occurred in each of the VCF files of the original HRC dataset). The excluded 100 samples were extracted from original HRC dataset to produce 100 separate VCF files.

The exact command line parameters are given below (VCF input dataset is `input.vcf`, compressed dataset is `archive`, input sample VCF is `sample.vcf` and archive of sample is `sample-arch`). The output was redirected to null device.

- Compression:

```
./gtshark compress-db -nl 20  input.vcf.gz archive
```

- Sample compression:

```
./gtshark compress-sample archive sample.vcf sample-arch
```

- Sample decompression:

```
./gtshark  decompress-sample archive sample-arch - > /dev/null
```

##### 4.4 Using population as a knowledge base

In this experiment, GTShark was used to compress VCF files describing single samples in reference to a previously compressed datasets being collections of samples from a single population or mix of populations.

The input data for this experiment (all input VCF files) was extracted from Chromosome 11 of the 1000GP3 dataset. It contains samples from 26 population, but populations ASW and MXL were excluded from test due to small number of related samples. The chosen populations are from 5 superpopulations: African (ACB, ESN, GWD, LWK, MSL, YRI), American (CLM, PEL, PUR), East Asian (CDX, CHB, CHS, JPT, KHV), European (CEU, FIN, GBR, IBS, TSI), South Asian (BEB, GIH, ITU, PJL, STU). Initially, 2040 samples from these 24 populations (85 samples from each population) were chosen for the test; a VCF file for each sample was extracted from the original data. Next, 48 reference datasets (VCF files) were created:

- 24 datasets with single populations, 85 samples from a single population in each,
- 24 larger datasets with mixed populations, 85 samples from each of 23 populations (all but one), that is 1955 samples, in each dataset.

The reference datasets were compressed and each of the 2040 samples was compressed in reference to each of the 48 archives.

The exact command line parameters are given below (VCF input dataset is `input.vcf`, compressed dataset is `archive`, input sample VCF is `sample.vcf`, archive of sample is `sample-arch` and decompressed VCF is `out-sample.vcf`).

- Compression:

```
./gtshark compress-db -nl 0  input.vcf.gz archive
```

- Sample compression:

```
./gtshark compress-sample archive sample.vcf sample-arch
```

#### 5 Environment

The computer used in tests was of the following configuration:

- 2 AMD Opteron 6348 CPUs, 12 cores per CPU, each clocked at 2.8 GHz,
- 128 GiB RAM.

For compilation we used G++ v. 6.3.0. The machine was running Debian GNU/Linux 9.5.

### 6 Additional results

Complete results are in the supplementary spreadsheet **“GTSharkEvaluation”**. This section provides summary of the results and points to the related full data in the spreadsheet.

Note that compressed sizes and archive sizes (in the main manuscript and in this document) refer to sizes of compressed genotypes only. Sizes of other components of archives (compressed description of variants and samples) can be found in the supplementary spreadsheet.

#### 6.1 Compression and decompression of a full dataset—compressors comparison

A summary comparison of compressors of VCF files for HRC collection is in Table 1 in the main manuscript. A similar comparison for 1000GP3 data is shown here in Table 1. Detailed results for both datasets, distinct for each of the input VCF file, are in the **“HRC dataset compression”** and **“1000GP3 dataset compression”** sheets of the supplementary spreadsheet. These spreadsheets contain some characteristics of the input VCF files (size, gzipped size, number of variants, number of samples) and, for each of the evaluated programs, complete sizes of compressed structures (genotypes and description of variants and samples), as well as times and memory usage of compression and decompression stages. The results are provided for all evaluated tools: TGC, PBWT, BGT, GTC, GTShark. In case of GTShark, the results are for different values of `nl` parameter.

Comments on the results:

- due to limitations of TGC (see Section 4.1), the TGC compression and decompression times refer to compressing and decompressing BV (Bit Vectors) only, time and memory usage of transforming VCF files to BV are in the supplementary spreadsheet,
- BGT results for HRC dataset are incomplete: VCF file of chromosome X_nonPAR_MALE could not be compressed, as BGT does not support haploid genotypes,
- BGT results for 1000GP3 dataset are incomplete: VCF file of chromosome Y could not be compressed, as BGT does not support haploid genotypes,
- PBWT results for 1000GP3 are incomplete, as program failed to compress VCF file for chromosome X.

Table 1: Comparison of compressors of VCF files for 1000GP3 collection (84.8 M variants, 2,504 samples, 854 GB of VCF, 17.4 GB of gzipped VCF).

| | C-size [MB] | C-time [s] | C-RAM [GB] | D-time [s] |
|---|---|---|---|---|
| TGC* | 852 | 42,339 | 11.5 | 449 |
| PBWT* | 1,460 | 25,186 | 1.0 | 5,229 |
| BGT* | 3,307 | 29,012 | **0.004** | 5,443 |
| GTC | 1,247 | **25,586** | 1.2 | **735** |
| GTShark | **649** | 28,421 | 4.1 | 4,655 |

Column description: ‘C-size’ — compressed size, ‘C-time’ — compression time, ‘D-time’ — decompression time, ‘C-RAM’ — amount of RAM in the compression stage. Bold font is used for the best results (TGC times are not taken into account, as they refer only to the compression and decompression of BV). * See comments on the results in Section 6.1.

#### 6.2 Extraction of a single sample

Extraction of a single sample from a GTShark compressed HRC dataset takes 691.83 seconds on average. Extraction of a single sample from a GTShark compressed 1000GP3 dataset takes 822.98 seconds on average.

Detailed results for both datasets, distinct for each of the input VCF file (chromosome), are in the **"Sample extraction"** sheet of the supplementary spreadsheet. The list of samples used in the experiment is also included.

#### 6.3 Compression and decompression of a single sample using previously compressed dataset as a reference

The compression of a single sample using previously compressed dataset as a reference (HRC dataset with 100 samples excluded) takes 715.56 seconds on average. The decompression takes 765.05 seconds on average. The size of the compressed sample is 65,501.48 B on average.

Detailed results, distinct for each of the input VCF file (chromosome), are in the **"Sample compression"** sheet of the supplementary spreadsheet. The list of samples used in the experiment is also included.

#### 6.4 Using population as a knowledge base

The main results are in Figure 1 in the main manuscript. A more detailed results are in the **"Population as a knowledge base"** sheet of the supplementary spreadsheet, which includes four tables:

- Table 1, which is a matrix of sizes of a samples compressed using a reference made of 85 samples from a single population (average over 85 samples); this is the matrix included in Figure 1 in the main manuscript,
- Table 2, which is a matrix of sizes of a samples compressed using a reference made of 1955 samples from 23 populations (average over 85 samples); the diagonal of this matrix is the additional column in Figure 1 in the main manuscript,
- Table 3, which presents sizes of the references used,
- Table 4, which presents per sample sizes of the references used.
